# Aquatic respiratory rates in red devil vampire crabs (*Geosesarma hagen*) are dependent on interactions between temperature, sex, and body size

**DOI:** 10.64898/2026.06.30.735571

**Authors:** Ginger A. Buck, Bryan H. Juarez, Madison P. Lacey, Lauren A. O’Connell, Victoria M. Watson-Zink

## Abstract

The shift to terrestrial environments in ancestrally aquatic animals is often associated with key physiological and physical changes, including shifts in respiratory physiology and in some cases, even the evolution of completely novel respiratory structures. Examining how respiration operates across a gradient of submersion states in ancestrally aquatic terrestrial animals may shed light on how complex biological traits shift under different selective regimes. In this work, we begin exploring respiration in terrestrially-adapted land crabs that still use their gills to respire while underwater. We tested the relationship between aquatic respiratory rates, body size, and sex in red devil vampire crabs (*Geosesarma hagen*) at two ecologically-relevant temperatures.

We found small females respire more than small males at 28°C, while large females respire more than large males at 21°C. Additionally, body size is a significant factor affecting respiratory rates of both sexes at 21°C and warmer temperatures significantly increase respiration in small crabs of both sexes. Interactions between these factors also led to emerging trends that can be explained by both physiological rules, such as reproductive investment and surface-to-volume ratios and heat transfer. We also report a temperature coefficient (Q_10_) of 1.52 for this species, showing an expected 52% change in respiratory and metabolic rate for every 10°C increase. This work also demonstrates the importance of understanding how and to what extent biological variables like sex and body size interact with abiotic environmental factors when measuring physiological traits in ectothermic invertebrate animals.

**What is already known:** As crabs colonize terrestrial environments, they undergo adaptive physical and physiological changes in their respiratory abilities. Few studies have examined these respiratory shifts in individual terrestrial crab species using modern respirometry tools and techniques and across different submersion states and biotic and abiotic factors. We quantify how temperature, body size, and sex affect aquatic respiratory rates in adult terrestrial vampire crabs.

**What this study adds:** We found small females respire more than small males at 28°C, while large females respire more than large males at 21°C. Additionally, body size is a significant factor affecting respiratory rates of both sexes at 21°C and warmer temperatures significantly increase respiration in small crabs of both sexes. Interactions between these factors also led to emerging trends that can be explained by both physiological rules, such as reproductive investment and surface-to-volume ratios and heat transfer. We also report a temperature coefficient (Q_10_) of 1.52 for this species, showing an expected 52% change in respiratory and metabolic rate for every 10°C increase.

## Introduction

The patterns of animal biodiversity we see on land today can be traced back to several convergent transitions from the marine to the terrestrial realm long ago. At least seven major animal lineages (e.g. millipedes, arachnids, insects, vertebrates, isopods, mollusks, crustaceans, etc.) colonized land between the late Cambrian and the mid-Cretaceous and these clades have given rise to nearly all the species that currently inhabit terrestrial environments (Boyce and Nelsen 2025). Ancestrally marine animals that have colonized terrestrial environments had to overcome significant physical and chemical challenges associated with shifts in gravity, water and salt availability, ultraviolet radiation, and temperature as they made these transitions (Webb 2012). These animals have therefore had to modify nearly every aspect of their physiology to maintain homeostasis on land. And indeed, sea-to-land transitions are typically associated with major physiological changes in osmoregulation and excretion, thermoregulation, respiration, reproduction, and development (Wei et al. 2026).

During the process of terrestrialization, key morphological and physiological traits evolved toward new adaptive optima (Lo Cascio Sætre et al. 2017). Some traits have been lost, as seen in the loss of the pelagic developmental stages in semi-terrestrial freshwater crayfish (Vogt and Tolley 2004), while other traits have acquired dramatic new functions in response to the novel selective pressures experienced on land, as seen in blennies that have evolved to respire effectively across their skin in the aerial supralittoral zone instead of their gills (Ord and Hundt 2020). Understanding how animals have repeatedly and convergently adapted to terrestrial environments, either through the co-option or loss of existing traits or the acquisition of new ones, is critical to understanding the history and origins of terrestrial life on Earth.

Decapod land crabs have convergently colonized land more than 18 times since the mid-Cretaceous, with more than 17 independent terrestrial transitions in the infraorder Brachyura (true crabs) and at least one in the infraorder Anomura (false crabs) (Wolfe et al. 2024). These transitions have led to many dramatic changes in the biology and physiology of terrestrial crabs (Watson-Zink 2021). In terms of respiratory adaptations, understanding how the ancestral trait of gill-driven aquatic respiration has shifted in land crabs is still an open question in the field.

Generally, there has been a progressive shift from a respiratory system driven by aquatic respiration via the gills to one reliant instead on aerial respiration via either a smooth or invaginated lung that lines the interior surface of the gill chambers (Taylor & Innes 1988; Diaz and Rodriguez 1977; Felgenhauer and Abele 1983; Morris 2002).

At one end of this respiration evolutionary pathway, some semi-terrestrial crabs still primarily respire with their gills, as in the intertidal green crab, *Carcinus maenas*. These crabs hold seawater inside of their branchial chambers, sometimes for up to six hours, and can withstand extreme hypercapnic conditions during low tide exposure periods (Simonik & Henry 2014). At the other end, some species are so terrestrially adapted that they breathe solely with their branchiostegal lungs and will drown if immersed for an extended period; in these taxa, the gills then take on a new role in osmoregulation and excretion, as in the coconut crab, *Birgus latro*, and the Christmas Island red crab, *Gecarcoidea natalis* (Greenaway et al. 1988; Morris 2002; Farrelly & Greenaway 1994). And many other species fall between these two extremes, as in the ghost crabs, *Ocypode sp*., and the estuarine crab, *Chasmagnathus granulatus*, which both display bimodal respiration. Bimodally respiring crabs respire with their gills when submerged, but are able to extract oxygen directly from the air via the branchiostegal lungs while emersed (Halperin et al. 2000; Al-wassia et al. 1989; Whiteley et al. 1990; Gannon and Henry 2004). In some clades, the respiratory shift from gills to lungs has also been associated with both reductions in the number of phyllobranchiate gill pairs and overall reductions in gill size in some clades (Burggren and McMahon 1988). Yet other groups have alternatively evolved tympana, which are membranous windows on the legs and thoracic sterna that assist in aerial respiration (Maitland 1986).

What the field lacks, however, are studies that rigorously explore respiration in land crab species that display varying degrees of terrestrial adaptation across a gradient of submersion states using modern respirometry. O’Mahoney and Full (1984) investigated how three different crab species with various degrees of terrestrial adaptation respired in both air and in water. They found that the terrestrial *Gecarcinus lateralis* had the highest oxygen consumption in air, while the aquatic *Callinectes sapidus* had the lowest oxygen consumption. Conversely, in water, this trend was flipped with the terrestrial crab having the lowest oxygen consumption while immersed, and the aquatic crab having the highest oxygen consumption. The “amphibious” crab, *Cardisoma guanhumi*, maintained a constant rate of oxygen consumption volume across both media. They explained their findings by suggesting that these differences in oxygen consumption across the three species resulted from differences in gill morphology, the presence/absence of lung tissue in the branchial chambers, and changes in the ventilation rates of the amphibious crab to accommodate differences in oxygen consumption in air and in water, which is a physiological compensation that was not observed in either the terrestrial or the aquatic crab while they were in their non-preferred respiratory medium. Finally, they reported that the terrestrial *G. lateralis* was only able to survive up to 18 hours underwater, while the aquatic *C. sapidus* could only survive for 24-48 hours in air (O’Mahoney and Full 1984).

More recently, in terms of a single species, Cumberlidge (1991) investigated the “respiratory formula” in terrestrial *Globonautes macropus*, measuring how respiration changed across a gradient of submersion states while also recording which tissues and structures were involved in respiration in each instance (Cumberlidge 1991). He found that *G. macropus* has reduced the surface area taken up by its gills to allow for a correspondingly larger area for its lung. It has also reduced its number of gill pairs from nine to seven by removing two of its anterior gill pairs, which is an adaptation that has also been seen in other land crabs that are adapted to extremely arid environments. This crab is also capable of respiring with its gills and its lungs simultaneously, as in *C. guanhumi*, bestowing it with the ability to survive in fully aquatic conditions in spite of its highly terrestrial nature (Cumberlidge 1991). Few land crab species have been studied this extensively, however, and even fewer using modern respirometry approaches, making it difficult to make comparisons across taxa, especially across clades with varying degrees of terrestrial adaptation.

In this work, we aim to begin filling this gap by exploring respiration in terrestrial vampire crabs (*Geosesarma sp*.) which, within a single genus, display wide variation in their relative degrees of terrestrial adaptation. Vampire crabs (*Geosesarma sp*.) are endemic to Java, Indonesia, where surface temperatures average 31°C and range from 16°C to 41°C (Sholihah and Shibata 2019; Jaelani and Handayani 2022). They are commonly found many kilometers from shore in muddy, wet burrows near freshwater streams. In terms of their respiratory capabilities, vampire crabs are thought to respire with their gills to some degree when submerged and with a novel respiratory tissue (the Verwey tissues) located on the inner surface of their pterygostomial region, which acts as a rudimentary lung when they are on land (Felgenhauer and Abele 1983), but their relative respiratory abilities across a gradient of submersion states has never before been studied. In captivity, while the crabs are usually found on land, albeit hidden in various cryptic locations in their enclosures (e.g. in shallow burrows in the substrate, beneath driftwood pieces underneath moss sheets, etc.), we have also observed them completely submerged for several hours at a time in the water dishes, implying that they possess some ability to withstand full submersion for moderate amounts of time, although not indefinitely.

In this exploratory work, we sought to establish a baseline for measuring respiration in direct-developing vampire crabs by first establishing how they breathe in ancestral aquatic conditions.

Previous studies have shown that sex (Shillington 2005), body size (Huebner 1973), and temperature (Bonacina et al. 2022) impact respiration rates in ectotherms. To gain a complete picture of respiratory physiology in this clade, in a similar fashion, we also measured respiratory rates in red devil vampire crabs (*Geosesarma hagen*) using both male and female crabs across a range of body sizes and water temperatures. We hypothesized that, due to the interactions between sex and reproductive capacity, body size, and ambient temperature, that large females in warmer temperatures would have the highest respiratory rates while small males in cooler temperatures would have the lowest respiratory rates. This work will lay the groundwork for future studies that seek to examine how their respiration shifts with their submersion status (i.e., partially submerged or completely out of water), and how these patterns track across congeneric species with differing degrees of terrestrial adaptation.

## Methods

### Animal Acquisition and Care

We obtained N = 15 adult intermolt red devil vampire crabs, *Geosesarma hagen* (Ng et al. 2015) from Aqua Imports (Boulder, CO, USA). Of the 15 crabs, seven were females, five of which were brooding. These animals were housed individually in transparent plastic containers (18.09 cm L x 11.13 cm W x 13.34 cm H) within opaque plastic containers with mesh-covered holes for air circulation and light penetration. We added cardboard barriers between each crab to minimize potential aggression. Each tank contained about five centimeters of moist soil mixed with sphagnum moss, a small cup for shelter, and another small cup for water with a shallow layer of mineral clay. We kept the animals in a humidity-controlled (80% relative humidity) room kept at an average of 26°C with a 12 hour light-dark cycle. Crabs were fed ¼ teaspoon of *Drosophila hydei* that were dusted with Repashy Calcium Plus (Repashy Ventures, Inc.; Oceanside, CA, USA) three times per week. Each crab was also given three pinhead crickets twice a week. All water for their care and experimentation was distilled water mixed with SaltyShrimp Mineral GH/KH+ Powder (Garnelenhaus, Germany). All animals were allowed to acclimate to lab housing conditions for at least 2 months prior to experimentation. We fasted all animals for 24 hours prior to experimentation.

### Data Collection

Before experimentation, we sexed each crab and measured its carapace dimensions and wet weight. We used carapace width as a measure of body size since it, unlike body mass, is not heavily influenced by the amounts of non-respiring biominerals and biopolymers in the carapace.

We used the Q-Box AQUA Aquatic Respirometry Package from Qubit Systems (Kingston, Ontario, Canada) to measure oxygen use and water temperature through time (Fig. S1). While we considered using intermittent-flow respirometry (Svendsen et al. 2016; Killen et al. 2021) to estimate respiration rates in *G. hagen*, doing so would require submerging the crabs for extended periods of time, which could have negatively impacted its health and survival since the crabs cannot remain submerged indefinitely without drowning. For a given total trial duration, there is also a trade-off between the number of cycles and the duration of the measurement period per cycle. On one hand, given total trial duration, increasing the number of cycles comes with the drawback of increasing the animal’s time spent underwater when no data is collected during the flush periods and wait periods that precede the measurement period. Relatively shorter measurement periods are also associated with lower signal-to-noise ratios (Svendsen et al. 2016). On the other hand, increasing the duration of the measurement period per cycle limits the total number of cycles (the sample size) obtained during each trial. Therefore, while intermittent-flow respirometry is an appropriate standard for most aquatic animals, it may not be suitable for all animals such as those that are terrestrial but ancestrally aquatic. Furthermore, high rates of mortality can result when submerging some terrestrial crab species in water for extended periods (Watson-Zink et al. 2024). Given the described limitations of working with our experimental animals using standard methodology, we used closed (stop-flow) respirometry to quantify respiration rates. Closed respirometry allowed us to reduce the total experimental period per animal while increasing time spent in the measurement period since only a single flush and wait period is required prior to measurement. While extended measurement periods in closed respirometry risks exposing the animals to anoxic conditions and can bias measurements (Steffensen 1989), our experimental design reduced this risk and we confirmed empirically that the animals did not experience anoxic conditions.

We follow the methods of Killen et al. (2021) in describing our respirometry setup and experimental protocol. We acclimated the animals in the measurement chamber for 10 minutes in flowing water. We recorded data during the measurement period for each crab for at least 16 minutes and we terminated trials at either 35 minutes or when the water reached 90% of the initial dissolved oxygen to minimize respiratory stress. The volume of the empty acrylic respirometry chamber connected with simple plastic tubing was 42.74 mL, the volume of the chamber alone was 7.07 mL, and the animal masses ranged from 0.70–1.59 g. We accounted for all tubing volume when measuring respiration. Chamber mixing was achieved using an in-line pump and a flow rate of 38 mL per minute (corresponding to a pump voltage of 0.95). We used an S122 Optical dissolved oxygen probe from Qubit Systems, which was placed in the mixing circuit, to measure dissolved oxygen. The dissolved oxygen, temperature, and pressure probes were calibrated by Qubit and we obtained all measurements within 3 months of purchasing the equipment. The Qubit software LoggerPro version 3.16.2 performed automatic temperature compensation during dissolved oxygen probe usage. We cleaned all equipment daily with 90% ethanol, including all tubing. Additionally, we cleaned the container holding water for the flushing reservoir with soap and water before rinsing it thoroughly, drying it, and cleaning it with 90% ethanol. Using the latter flushing reservoir, we aerated 3.25 L of SaltyShrimp-treated water overnight at 21°C or 28°C.

We randomly assigned crabs to each temperature treatment and placed them in aerated water at either 21°C or 28°C. These temperatures were chosen as a compromise between minimizing discomfort in the crabs and covering a wide range of temperatures experienced by the species in nature (Sholihah and Shibata 2019). The experiment was run inside a Fisherbrand Isotemp incubator (Santa Clara, California, United States) to stabilize water temperatures. We obtained respiratory measurements during the day in dim light to further minimize stress by covering an LED light inside the incubator with thin cardboard paper. We measured 15 crabs, one at a time, over 8 total days (1 or 2 animals per day). During measurement, the animals were shielded from visual stimuli by the chamber, the chamber’s holding container, and the incubator.

### Data Analysis

We standardized all measurements to a sampling frequency of 12 min^−1^ (sampling every 5 seconds). We selected this sampling frequency as a trade-off between sampling often and reducing noise in the data. We subsetted the data to the sampled time shared across all crabs (16 minutes). Following standard protocol, we filtered out the first 25% of the data to reduce noise associated with starting each trial (Qubit Systems Inc., 2022), leaving 12 min (=144 samples) per individual. Next, we estimated the amount of oxygen in the respirometer using Eqn 1.

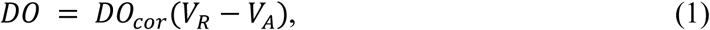

where DO is dissolved oxygen (mg O_2_), DO_cor_ is the per L amount of oxygen in the respirometer after correcting for salinity (mg O_2_ L^−1^), V_R_ is the respirometer volume (L), and V_a_ is the animal volume. We selected an *a priori* cutoff *r^2^* of 0.95 (Svendsen et al. 2016) relating DO and time, but removed no samples since the minimum r^2^ was 0.988.

We analyzed these data by fitting a generalized linear mixed model using nlme version 3.1-168 (Pinheiro et al., 2025; Pinheiro and Bates, 2000) in R version 4.5.0 (R Core Team, 2024). We regressed DO onto the main effects and interactions among time, temperature, body mass, and sex, and included crab ID as a random effect. The latter procedure quantifies respiratory rates in units of mg O_2_ min^−1^ across combinations of temperature, body mass, and sex.

Furthermore, analyzing repeated measurements of DO through time violates the regression assumption of independent samples since the DO samples are autocorrelated through time. We accounted for this temporal autocorrelation by including an autoregressive and moving average (corARMA) correlation structure for each ID (Pinheiro and Bates 2000; Box et al. 2015). We co-optimized the autoregressive (*p*) and moving average (*q*) parameters by varying each between 0 and 5, and comparing fits using AICc, the Akaike Information Criterion corrected for small sample size (Hurvich and Tsai 1989). For interpretation, we selected the best model with the lowest AICc and no other model was within 2 AICc units of the best model.

We identified significant interactions among discrete and continuous variables in the best model. Interactive effects are typically determined using estimated marginal (least-squares) means across the range of independent variables (factor levels or extremes of a continuous range). Interactions among discrete and continuous variables require comparing slope effects across the levels of the discrete factor (Searle et al. 1980). Thus, our post-hoc analysis involved comparing the estimated marginal means of the respiration rate (slope effect) across the temperature treatments (21 and 28°C), body size, and sex using the emtrends function in the emmeans library version 1.11.0 (Lenth, 2023). We used a conservative range of carapace widths (the 1st and 3rd quartile; 11.43 and 12.95 mm) to estimate marginal means instead of the maximum and minimum to avoid modeling unrealistic body sizes for each sex since the species is sexually dimorphic in size (males tend to be larger than females). We used two statistically conservative techniques when comparing respiration rates across temperature, body size, and sex. First, instead of performing all possible pairwise comparisons (N = 28) across 8 groups (all combinations of two temperatures, two sizes, and two sexes), we created contrasts to compare the marginal effects of one variable while holding the other two constant. The latter resulted in N = 12 total pairwise comparisons. For example, we tested for an effect of temperature on the respiration rate while holding mass and sex constant. Second, we adjusted for multiple comparisons using the conservative Bonferroni correction. This methodology reduces experiment-wide Type I error rates. Finally, we regressed size-specific respiratory rate against temperature using ID as a random effect and used the model coefficients to estimate the respiratory Q_10_ for *Geosesarma hagen*, a metric that reports how much an animal’s respiratory rate increases per 10°C increase in temperature.

## Results

We found that temperature, body mass, and sex had non-additive effects on respiratory rates (Table 1; Table S1). While respiratory rates seemed to increase with greater temperatures, larger body sizes, and in females relative to males (Fig. 1), as we predicted, the significant interaction between time, temperature, body mass, and sex indicated that interpreting the combined effects of each variable is necessary. Thus, we reported marginal effects (the respiratory rate) and confidence intervals for each combination of traits (Table S2). However, we interpreted only the 12 (of a possible 28) pairwise comparisons which held two of the three covariates constant (Table 2).

**Figure 1.**
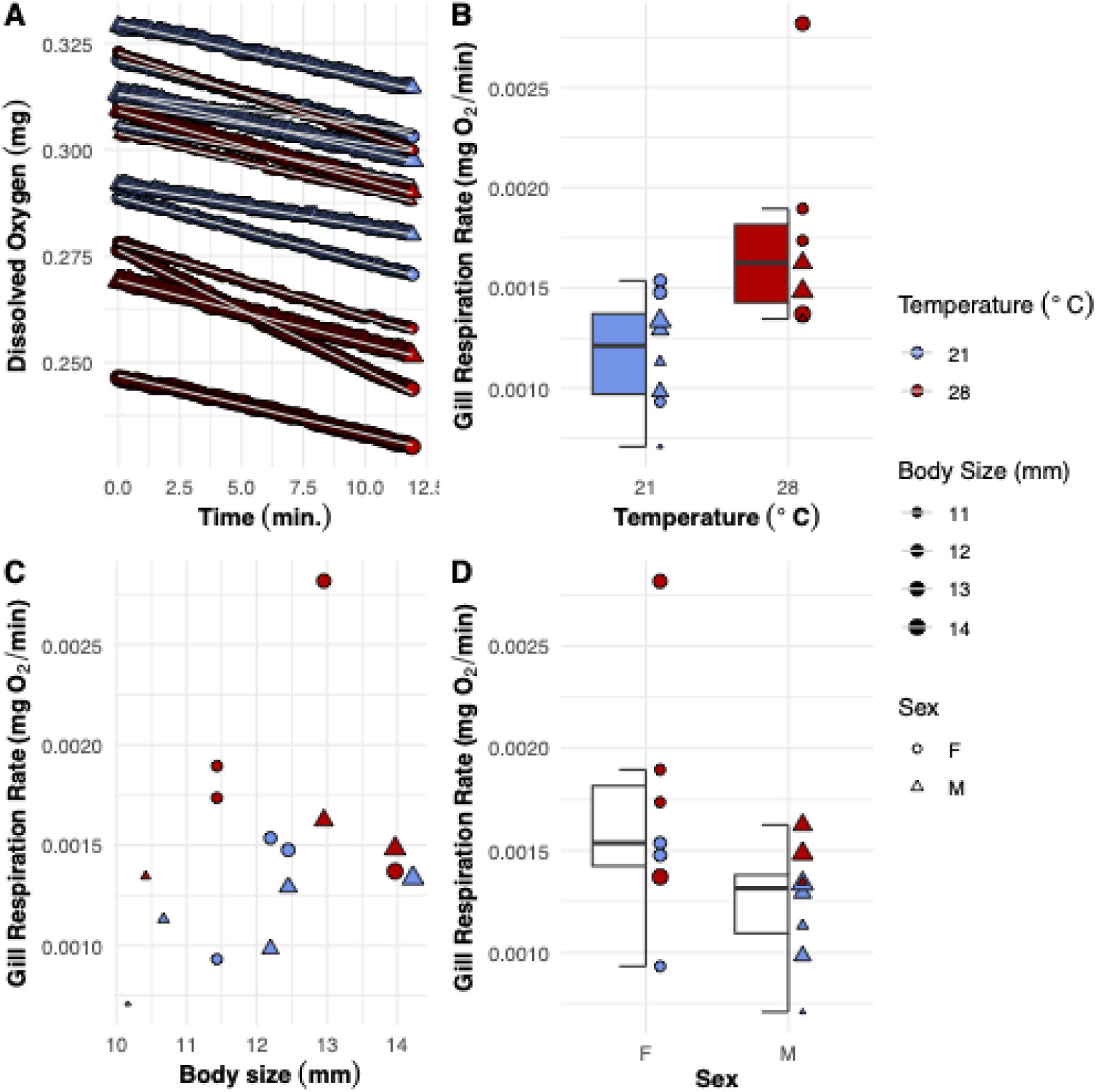
Structure of the raw data on respiratory rate. The shared legend denotes 21°C as blue and 28°C as yellow, larger animals as larger symbols, and males as triangles and females as circles. **A.** Measured dissolved oxygen through time. **B.** Measured respiratory rates at each experimental temperature. **C.** Measured respiratory rates across body size. **D.** Measured respiratory rates for each sex.

**Table 1.** The best fit generalized linear mixed model of the effects of time, temperature, body size, and sex on Dissolved Oxygen (DO). The selected autoregressive and moving average parameters were p = 4 and q = 0. Temp = temperature, Size = carapace width, M = male, β = regression coefficient, SE = standard error, DF = degrees of freedom. Coefficient units are in parentheses. Rows with significant p-values are in bold.

| Model Term | $\beta$ | SE | DF | t-value | p-value |
| --- | --- | --- | --- | --- | --- |
| Intercept (mg O<sub>2</sub>) | -5.33e-01 | 8.32e-01 | 2133 | -0.641 | 0.522 |
| **Time (mg O<sub>2</sub> min<sup>-1</sup>)** | **-4.52e-02** | **1.13e-02** | **2133** | **-3.985** | **<0.001** |
| Temp (mg O<sub>2</sub> °C<sup>-1</sup>) | -1.03e-04 | 3.15e-02 | 2133 | -0.003 | 0.997 |
| Size (mg O<sub>2</sub> mm<sup>-1</sup>) | 1.65e-02 | 6.96e-02 | 11 | 0.237 | 0.817 |
| M (mg O<sub>2</sub>) | 4.17e-01 | 8.60e-01 | 11 | 0.485 | 0.638 |
| **Time:Temp (mg O<sub>2</sub> min<sup>-1</sup>°C<sup>-1</sup>)** | **1.75e-03** | **4.31e-04** | **2133** | **4.070** | **<0.001** |
| **Time:Size (mg O<sub>2</sub> min<sup>-1</sup>mm<sup>-1</sup>)** | **3.73e-03** | **9.47e-04** | **2133** | **3.935** | **<0.001** |
| Temp:Size (mg O<sub>2</sub> °C<sup>-1</sup>mm<sup>-1</sup>) | 1.38e-04 | 2.63e-03 | 2133 | 0.052 | 0.958 |
| **Time:M (mg O<sub>2</sub> min<sup>-1</sup>)** | **4.09e-02** | **1.17e-02** | **2133** | **3.498** | **0.001** |
| Temp:M (mg O<sub>2</sub> °C<sup>-1</sup>) | -9.30e-03 | 3.27e-02 | 2133 | -0.284 | 0.777 |
| Width:M (mg O<sub>2</sub> mm<sup>-1</sup>) | -3.32e-02 | 7.17e-02 | 11 | -0.462 | 0.653 |
| **Time:Temp:Size (mg O<sub>2</sub> min<sup>-1</sup>°C<sup>-1</sup>mm<sup>-1</sup>)** | **-1.39e-04** | **3.60e-05** | **2133** | **-3.861** | **<0.001** |
| **Time:Temp:M (mg O<sub>2</sub> min<sup>-1</sup>°C<sup>-1</sup>)** | **-1.56e-03** | **4.47e-04** | **2133** | **-3.500** | **0.001** |
| **Time:Size:M (mg O<sub>2</sub> min<sup>-1</sup>mm<sup>-1</sup>)** | **-3.37e-03** | **9.75e-04** | **2133** | **-3.457** | **0.001** |
| Temp:Size:M (mg O<sub>2</sub> °C<sup>-1</sup>mm<sup>-1</sup>) | 6.86e-04 | 2.73e-03 | 2133 | 0.251 | 0.802 |
| **Time:Temp:Size:M (mg O<sub>2</sub> min<sup>-1</sup>°C<sup>-1</sup>g<sup>-1</sup>)** | **1.28e-04** | **3.73e-05** | **2133** | **3.425** | **0.001** |

**Table 2.** Pairwise comparisons of respiration rates (mg O_2_ min^−1^) across temperature, body mass, and sex. Each contrast holds 2 of 3 variables constant; “at” indicates the margin over which comparisons were made. SF = small female, LF = large female, SM = small male, LM = large male, SE = standard error, DF = degrees of freedom, p-adj = Bonferroni adjusted p-value. The selected body masses, 0.82–1.08 g, are the range of body masses shared by both sexes; S = 0.82 g, L = 1.08 g. Rows with significant differences in metabolic rate are in bold.

| Contrast | Estimate | SE | DF | t-ratio | p-adj |
| --- | --- | --- | --- | --- | --- |
| **28°C v. 21°C at SF** | **1.15e-03** | **1.84e-04** | **2133** | **6.255** | **<0.001** |
| 28°C v. 21°C at LF | -3.34e-04 | 2.83e-04 | 2133 | -1.178 | 1 |
| LF v. SF at 28°C | -2.49e-04 | 1.32e-04 | 2133 | -1.882 | 0.719 |
| **LF v. SF at 21°C** | **1.23e-03** | **3.03e-04** | **2133** | **4.068** | **<0.001** |
| **28°C v. 21°C at SM** | **4.15e-04** | **1.20e-04** | **2133** | **3.447** | **0.007** |
| 28°C v. 21°C at LM | 2.94e-04 | 1.17e-04 | 2133 | 2.511 | 0.145 |
| LM v. SM at 28°C | 6.33e-05 | 8.42e-05 | 2133 | 0.753 | 1 |
| **LM v. SM at 21°C** | **1.83e-04** | **6.29e-05** | **2133** | **2.918** | **0.043** |
| LM v. LF at 28°C | -2.54e-04 | 1.45e-04 | 2133 | -1.752 | 1.000 |
| **LM v. LF at 21°C** | **-8.82e-04** | **2.17e-04** | **2133** | **-4.065** | **<0.001** |
| **SM v. SF at 28°C** | **-5.67e-04** | **1.39e-04** | **2133** | **-4.064** | **<0.001** |
| SM v. SF at 21°C | 1.68e-04 | 1.60e-04 | 2133 | 1.044 | 1 |

Generally, larger females at 28°C had the greatest average respiratory rate while smaller crabs of both sexes at 21°C had the lowest respiratory rates (Fig. 2). We also found that, on average, higher temperatures only significantly increased respiratory rates in small crabs of both sexes. At 21°C, we noted a definite impact of body size apparent across both sexes: larger female crabs had higher respiratory rates than smaller female crabs (1.23×10^−3^ mg O_2_ / min; t-ratio = 4.068, p_adj_ <0.001), and larger males had higher respiratory rates than smaller males (1.83×10^−4^ mg O_2_ / min; t-ratio = 2.918, p_adj_ = 0.043). We found sexual dimorphism in respiratory rates between small males and small females at 28°C (−5.67×10^−4^ mg O_2_ / min; t-ratio = -4.064, p_adj_ <0.001), and also between large males and large females at 21°C (−8.82×10^−4^ mg O_2_ / min]; t-ratio = - 4.065, p_adj_ <0.001). There were also temperature effects at small body sizes, but not large body sizes, for females (1.15×10^−3^ mg O_2_ / min]; t-ratio = 6.255, p_adj_ <0.001) and males (4.15×10^−4^ mg O_2_ / min]; t-ratio = 3.447, p_adj_ = 0.007). Finally, we report a respiratory Q_10_ for *Geosesarma hagen* of 1.520 over the temperature range of 21-28°C, indicating that the respiratory rate of this species increases by 52% for every 10°C increase in temperature.

**Figure 2.**
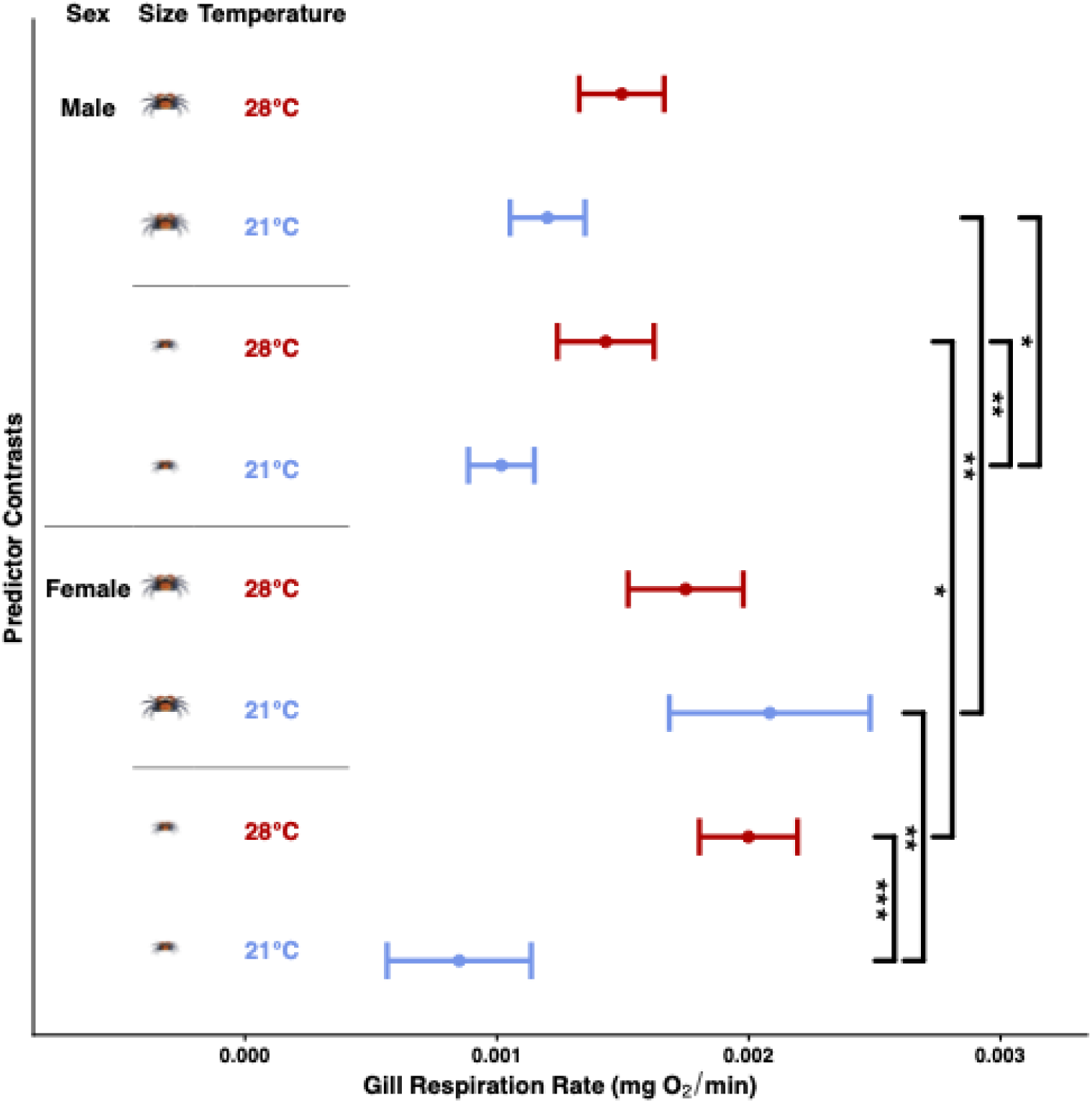
Pairwise slope tests of respiratory rate across combinations of temperature, body mass, and sex. Each pairwise comparison (predictor contrast) holds 2 of 3 variables constant and estimates are based on the best fit model. The pairwise slope comparisons tested are described in the main text and found in Table 2. Small and large body sizes correspond to the 25th and 75th percentiles of carapace widths (11.43mm and 12.954mm). Error bars indicate 95% confidence intervals. Significance levels (*** = 0.001, * = 0.05) correspond to significant respiratory rate differences.

## Discussion

The goal of this study was to understand how terrestrial vampire crabs (*Geosesarma*) respire in their ancestral aquatic habitat by examining their respiratory capabilities across both sexes, and a range of body sizes, and at two different ecologically relevant temperatures. Overall, we calculated that the Q_10_ of *G. hagen* (1.52) is greater than the average aquatic crab species which has a Q_10_ of 1.26 (Griffen and Sipos 2018). This indicates that while *G. hagen* is a primarily terrestrial species, as an ectothermic animal, its respiratory rate is still quite sensitive to temperature. Since Griffen and Sipos (2018) also described a link between metabolic rate and diet strategy, implying that higher metabolic rates are seen in carnivores, herbivores, and omnivores than in deposit feeders and detritivores due to variation in digestive structures, the higher Q_10_ in *G. hagen* may also be reflective of observed shifts in its diet towards carnivory and omnivory relative to the detritivorous diets of most aquatic crabs (Hohle and Singheiser 2016).

We also found that, instead of observing trends that would imply that our variables individually affect respiration (for example, females respire more than males, respiratory rates are greater at warmer temperatures), we found significant non-additive effects (interactions) between the variables, which we address individually below.

First, we found support for sexual dimorphism in respiratory rates in small crabs at 28°C and large crabs at 21°C (i.e. small females respire more than small males at 28°C, while large females respire more than large males at 21°C). The relative differences in reproductive investment due to sex (i.e. reproduction is more energetically costly for females than males; Smith et al. 2022) may explain this finding. To this point, we observed that five females in the experiment were gravid, carrying eggs at various stages of development. Because the metabolic state of the mothers changes when they are carrying eggs (Taylor and Leelapiyanart 2001) and the eggs themselves also respire (Simoni et al. 2011; Glazier 1991), we also anticipate that this may have affected our findings. There were not enough gravid females in this present study to test this effect statistically, but future research could explore how gravidity and egg-laying affect respiration rates in female crabs of varying sizes.

We also found that body size is a significant factor affecting respiratory rates of both sexes at 21°C (i.e. large crabs respire more than small crabs at 21°C regardless of sex). This finding is most likely the result of the combined effects of smaller crabs being more sensitive to warmer temperatures and having greater rates of heat transfer than larger crabs (Vandervoort 2020), and also smaller crabs having smaller gills than larger crabs, which would lead to differences in the rates of oxygen uptake resulting from differences in gill surface area-to-volume ratios. To further support the idea that smaller crabs are more sensitive to temperature changes than larger ones, we also calculated a Q_10_ of 2.38 for small crabs and a Q_10_ of 0.98 for large crabs, demonstrating that smaller animals are significantly more susceptible to respiratory changes at lower temperatures than larger crabs. Relatedly, we also found that warmer temperatures significantly increase respiration in small crabs of both sexes (i.e. small crabs respire more at 28°C than they do at 21°C, regardless of sex). Even in animals as distantly related as unicellular organisms and mammals, both body size and temperature have been found to be primary determinants for metabolic rates (and therefore, respiratory rates) (White et al. 2006; Gillooly et al. 2001; Clarke and Fraser 2004; Bihun et al. 2024), so our findings are consistent with patterns recorded in the literature.

Since vampire crabs are also capable of aerial respiration, future studies can also use the measured Q_10_ for aquatic respiration in *G. hagen* from this work to comparatively examine respiratory rates in vampire crabs across immersion states (i.e. emersed and partially submerged). This particular direction will be especially interesting because, while all marine ectotherms display a monotonic positive non-linear relationship between resting metabolic (respiratory) rate and temperature (Clarke and Fraser 2004), Addo-Bediako et al. (2002) found in a large-scale perspective work that across 346 different terrestrial insect species, there was a weak but significant negative relationship between respiratory rates and habitat temperatures.

This surprising result perhaps reflects fundamental differences in the physiology of marine ectotherms compared to terrestrial ones, or differences in thermal environment complexity that insects face on land (Clarke and Fraser 2004). Measuring if and how the relationship between respiratory rates and habitat temperatures shifts in vampire crabs when they are breathing aerially instead of aquatically may contribute critical data to our understanding of this phenomenon.

Finally, in terms of understanding the relationship between degree of terrestriality and respiratory abilities across a variety of submersion states, future studies should also include additional taxa that display differing degrees of terrestrial adaptation.

## List of symbols and abbreviations

DO: dissolved oxygen (mg O_2_)
DO_cor_: salinity-corrected oxygen density (mg O_2_ L^−1^)
V_R_: respirometer volume (L)
V_A_: animal volume (L).
GLMM: generalized linear mixed model
ARMA: autoregressive and moving average correlation structure
AIC: Akaike Information Criterion

## Acknowledgements

We thank the O’Connell lab at Stanford University for helping us care for the animals and providing valuable feedback on previous versions of this manuscript. We also thank Monika Kuzma and Billie Kearns from Qubit Systems Inc. for aiding us with the methodology.

This material is based upon work supported by the National Science Foundation Postdoctoral Research Fellowships in Biology Program under Grant No. 2109850 (to BHJ) and Grant 2109869 (to VWZ). VWZ was also concurrently supported by a Stanford Science Fellowship. GAB was supported by the Biology Summer Undergraduate Research Program (BSURP). LAO is a New York Stem Cell – Robertson Investigator.

## Author contributions

GAB: Conceptualization, Methodology, Investigation, Data curation, Writing (original draft), Writing (review and editing). BHJ: Conceptualization, Methodology, Software, Validation, Formal analysis, Investigation, Data curation, Writing (original draft), Writing (review and editing), Visualization, Supervision, Project administration, Funding acquisition. MPL: Investigation, Writing (review and editing). LAO: Conceptualization, Methodology, Resources, Writing (review and editing), Supervision, Project administration, Funding acquisition. VWZ: Conceptualization, Writing (original draft), Writing (final draft), Writing (review and editing), Supervision, Funding acquisition.

## Data availability

Data and code may be found on DRYAD (DOI: TO BE UPDATED UPON ACCEPTANCE).

## Supplementary Information

**Table S1.**
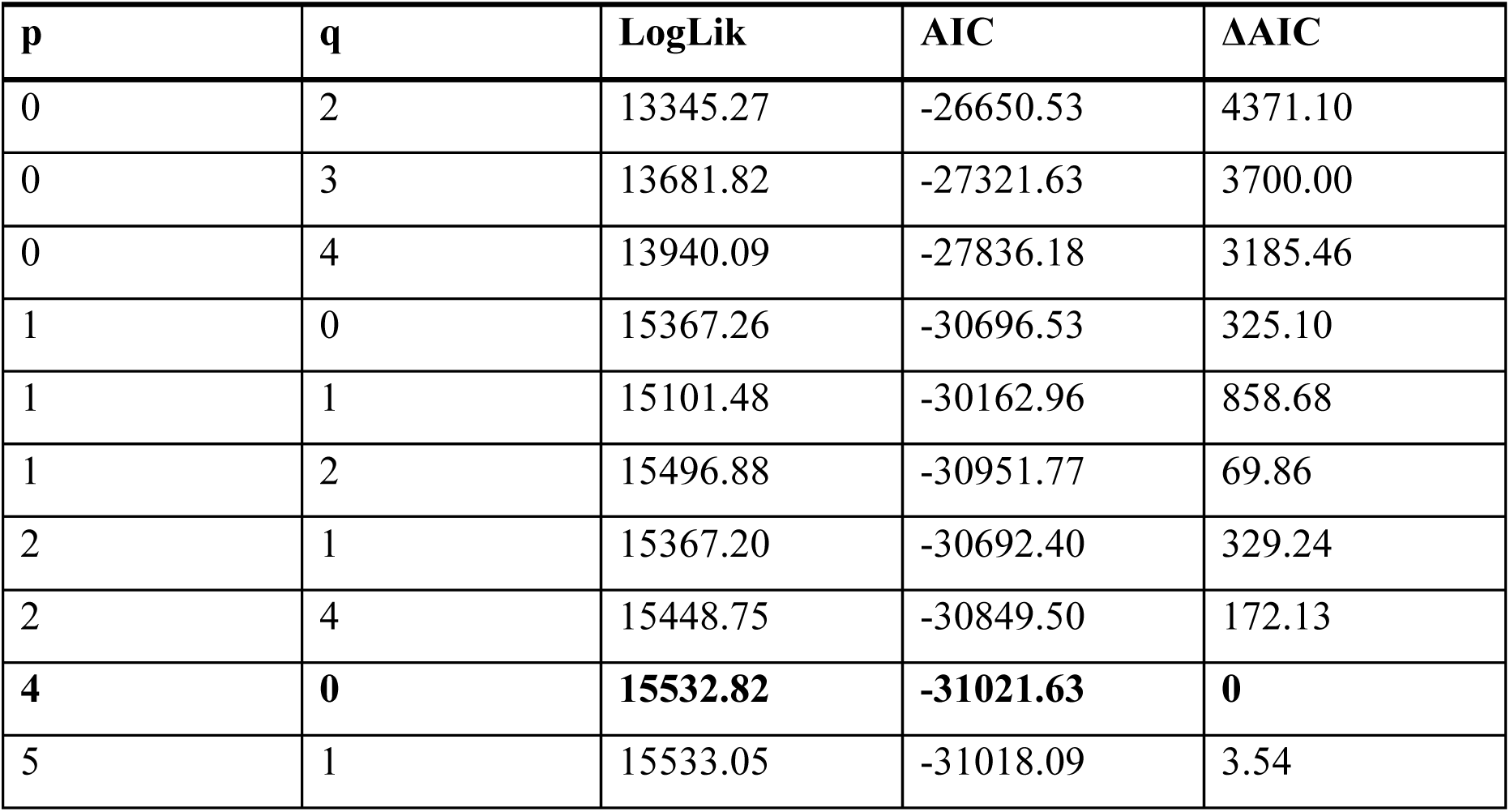
Model comparisons of temporal autocorrelation (ARMA) structures. P = autoregressive parameter, q = moving average parameter, LogLik = log-likelihood, AIC = Akaike Information Criterion, ΔAIC = AIC difference from the best fitting model. Models within 2 AIC units of the best model are in bold. Models for missing parameter combinations (p = 0–5; q = 0–5) did not converge.

**Table S2.** Marginal effects of time on dissolved oxygen (DO) across temperature, body size, and sex. Temp = temperature, Size = carapace width, SE = standard error, DF = degrees of freedom, CL = confidence level. The chosen margins for size are the 25th and 75th quantiles of carapace width.

| Temp | Size | Sex | Time Effect | SE | DF | Lower CL | Upper CL |
| --- | --- | --- | --- | --- | --- | --- | --- |
| 21 | 11.43 | F | 8.50e-04 | 1.46e-04 | 2130 | 5.64e-04 | 1.14e-03 |
| 28 | 11.43 | F | 2.00e-03 | 9.94e-05 | 2130 | 1.80e-03 | 2.19e-03 |
| 21 | 12.95 | F | 2.08e-03 | 2.03e-04 | 2130 | 1.68e-03 | 2.48e-03 |
| 28 | 12.95 | F | 1.75e-03 | 1.17e-04 | 2130 | 1.52e-03 | 1.98e-03 |
| 21 | 11.43 | M | 1.02e-03 | 6.69e-05 | 2130 | 8.87e-04 | 1.15e-03 |
| 28 | 11.43 | M | 1.43e-03 | 9.78e-05 | 2130 | 1.24e-03 | 1.62e-03 |
| 21 | 12.95 | M | 1.20e-03 | 7.60e-05 | 2130 | 1.35e-03 | 1.35e-03 |
| 28 | 12.95 | M | 1.50e-03 | 8.61e-05 | 2130 | 1.33e-03 | 1.66e-03 |

**Figure S1.**
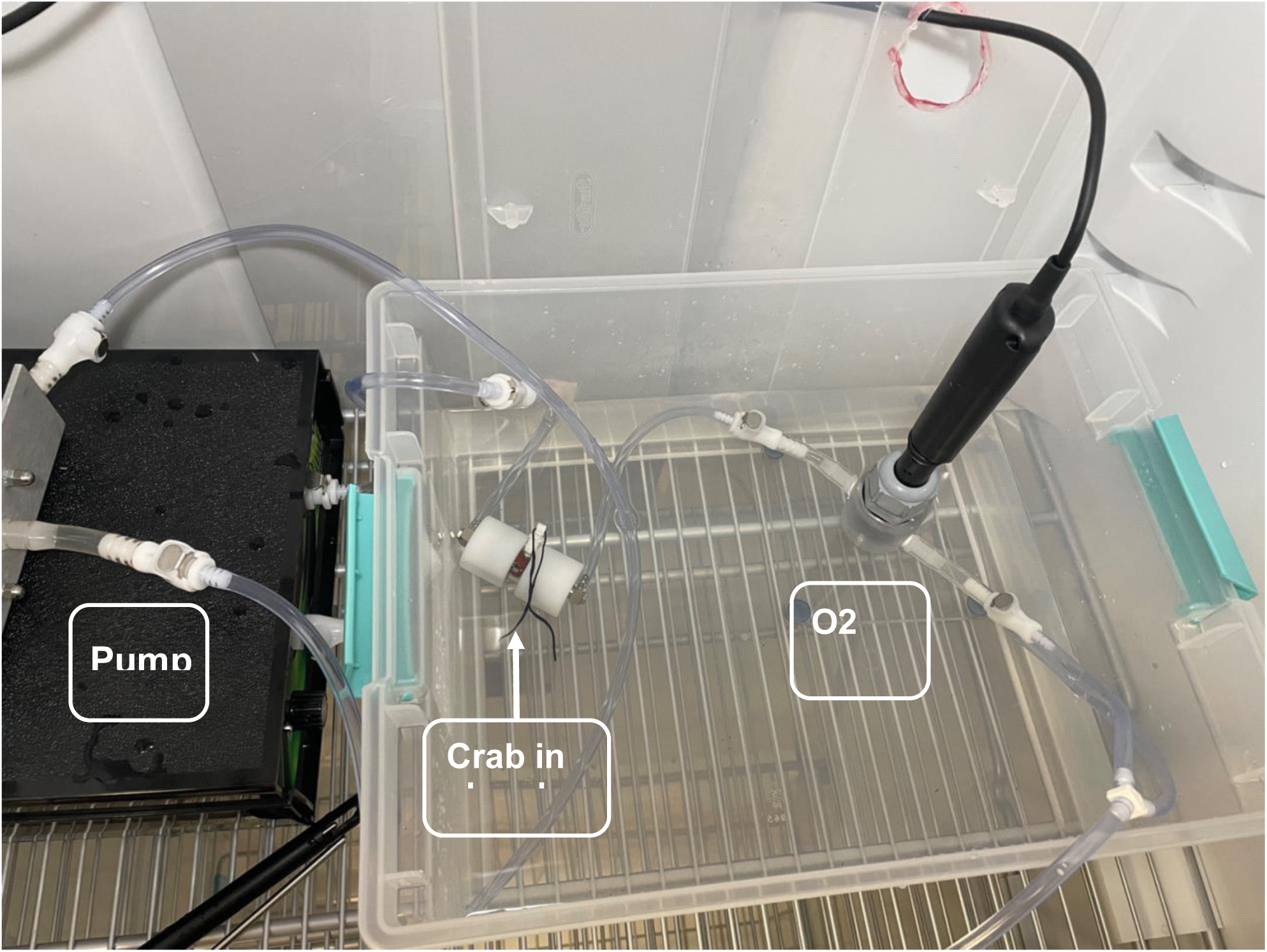
Photograph of respirometer setup. A crab is inside the animal chamber (with a black twist tie). The animal chamber is submerged in water. Respirometer is inside an incubator.

